# Universal nature of collapsibility in the context of protein folding and evolution

**DOI:** 10.1101/461046

**Authors:** D. Thirumalai, Himadri S. Samanta, Hiranmay Maity, Govardhan Reddy

## Abstract

Theory and simulations predicted sometime ago that the sizes of unfolded states of globular proteins should decrease continuously as the denaturant concentration is shifted from a high to a low value. However, small angle X-ray scattering (SAXS) data were used to assert the opposite, while interpretation of single molecule Forster resonance energy transfer experiments (FRET) supported the theoretical predictions. The disagreement between the two experiments is the SAXS-FRET controversy. By harnessing recent advances in SAXS and FRET experiments and setting these findings in the context of a general theory and simulations, we establish that compaction of unfolded states is universal. The theory also predicts that proteins rich in *β*-sheets are more collapsible than *α*-helical proteins. Because the extent of compaction is small, experiments have to be accurate and their interpretations should be as model free as possible. Theory also suggests that collapsibility itself could be a physical restriction on the evolution of foldable sequences, and provides a physical basis for the origin of multi-domain proteins.

## 1. PROTEIN COLLAPSIBILITY – WHAT IS THE PROBLEM?

The number of protein sequences with *N* amino acids that can be synthesized from twenty amino acids is 20^*N*^, which is approximately 10^130^ for *N* = 100. On the other hand, the number of folds in natural globular proteins, which may be associated with low energy compact structures, is only on the order of a few thousands [1, 2]. Clearly, the sequence space is dense in contrast to the sparse structure space. The dramatic reduction that occurs from the dense sequence space to the countable number of folds may be rationalized by merely imposing the restriction that folded globular proteins be Minimum Energy Compact Structures (MECS) [3]. Precise calculations using lattice models [4, 5] for proteins show that the number of compact structures grows exponentially with *N* [3], as predicted by polymer theory [6]. Remarkably, the number of MECS likely grows only as ln *N*, which has remained a surprising but under appreciated result [3]. The implication of this finding is that for a vast number of sequences the MECS must be topologically similar. In other words, the basins of attraction in the structure space are so rare that a vast number of sequences map on to precisely one structure with a specific topology. This plausibility, also established using lattice models [7], tidily explains the emergence of vastly limited number of structures from the astronomically large number of sequences. Thus, the propensity to form MECS is the distinguishing feature of biologically foldable sequences. Similar arguments could be made for RNA, which are made from four nucleotides. Indeed, it has been suggested that the requirement of compactness may be the key constraint for single stranded viral RNA evolution [8]. It is likely that compaction as a selection mechanism holds more generally for ribozymes as well.

The structures and dynamics of MECS as well the unfolded states are usually investigated by varying the concentration of denaturants, such as Urea or Guanidinium Chloride (GdmCl). A fundamental question that goes to the heart of protein collapsibility problem is: how do the shapes of the folded and unfolded states change as a function of the concentration of denaturants? In order to unpack the answer to this seemingly simple question, let us consider the folding of simple two-state proteins in which only the folded (*F*) and unfolded (*U_D_*) states are appreciably populated. The balance between the population of these *F* and *U_D_* states in experiments are altered by changing denaturant concentrations. The radii of gyration of the folded states, 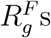, change imperceptibly [9] as the concentra-tion of denaturants, [*C*], is altered. However, as [*C*] decreases below the mid-point [*C_m_*], the concentration at which the folded and unfolded protein populations are equal, whether or not the size of the unfolded states, 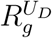, decreases becoming more compact has, until recently, remained in dispute [10]. In a nutshell, does the *U_D_* become compact forming the *U_C_* state, with 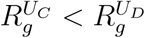 below [*C_m_*]?

Twenty five years ago, the answer to this question posed above was given in the affirmative [11] using theory. Subsequently by taking into account consequences of the finite size of globular proteins it was shown that folding cooperativity increases universally as ∼ *N*^2.2^, where *N* is the number of amino acids [12]. In the process, we argued that water is only a moderately good solvent, and is more likely to be closer to the solvent [12] (Box 1 gives a background of polymer physics terminology commonly used to analyze experimental data). However, analyses of the data using two different experimental methods have arrived at contradictory conclusions. Based on a number of Small Angle X-Ray Scattering (SAXS) experiments, it had been asserted emphatically for nearly two decades [10, 13, 14] that the dimension of the *U_D_* state does not change as [*C*] decreases. It, therefore, follows that 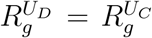 at all [*C*]. In sharp contrast, using single molecule Forster resonance energy transfer experiments (FRET), it was concluded that 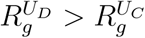 at low [*C*] [15–17], in accord with our theoretical predictions [3]. In light of experimental and theoretical advances in the last two years, we survey the current status and come to the conclusion that denatured state ensemble (DSE) collapse of single domain globular proteins is universal. As a corollary, we also suggest that, as a rule, Intrinsically Disordered Proteins (IDPs) must expand as the denaturant concentration increases.

#### Box 1. Polymer physics language for describing states of proteins

Despite substantial differences between proteins and homopolymers, the language and concepts to describe the latter, principally developed by Flory [18], have been adopted to characterize *U_D_* states and IDPs. Folded states of globular proteins are roughly spherical and are nearly maximally compact with high packing densities [19–21]. The radius of gyration 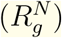 of folded proteins is well described by the Flory law with 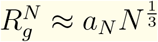, with *a_N_* ≈ 3.3 Å [22]. At high denaturant concentrations, proteins swell adopting expanded conformations. In unfolded *U_D_* states 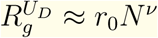 where *v* ≈ 0.6 is the Flory exponent with *r*_0_ ≈ 2.0 Å [23]. Estimates of *a_D_* vary greatly and is one of the difficulties in assessing solvent quality (Box 2). Thus, viewed from this perspective, we could surmise that proteins must undergo a coil-to-globule transition [24–26], a process that is reminiscent of the equilibrium collapse transition in homopolymers with *N* ≫ 1 [27, 28]. The latter is driven by a balance between conformational entropy and intra-polymer interaction energy. By analogy, we surmise that the swollen state is realized in good solvents (interaction between proteins and solvents is favorable) whereas in the collapsed state intra protein interactions are preferred. It is tempting to identify high (low) denaturant concentrations with good (poor) solvent for proteins. The simple physical picture given above is not wholly accurate because two additional states need to be considered in order to understand the collapsibility problem in polypeptide chains. First, upon increasing the denaturant concentration from zero, the side chains, which are densely packed in the *F* state, could become disordered while preserving the overall fold. Such a state is referred to as the dry globule (*DG*). Experiments [29, 30], theory [31] and simulations [32, 33] have provided evidence for the *DG* state, which we postulated to be a universal intermediate [34] in the folding landscape (Figure 1). As the denaturant concentration decreases, the *U_D_* state becomes compact, forming the *U_C_* state. These states (*F*, *U_D_*, *DG*, and *U_C_*) can be distinguished using two order parameters. One is the density, 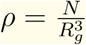 and the other is *χ*, which measures how similar a given conformation is to the *F* state [35]. If the protein is folded then 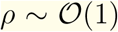 and *χ* ∼ 0 whereas in the *U_C_* state the value of *ρ* are small and 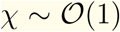. Similarly, the *DG* state is characterized by 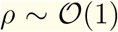 and *χ* ≠ 0, but not too large. Finally, the value of *ρ* in *U_C_* is greater than in the *U_D_* state whereas *χ* is smaller. Figure 1 illustrates the states of a globular protein as a function of *ρ* and *χ*.

**FIG 1.**
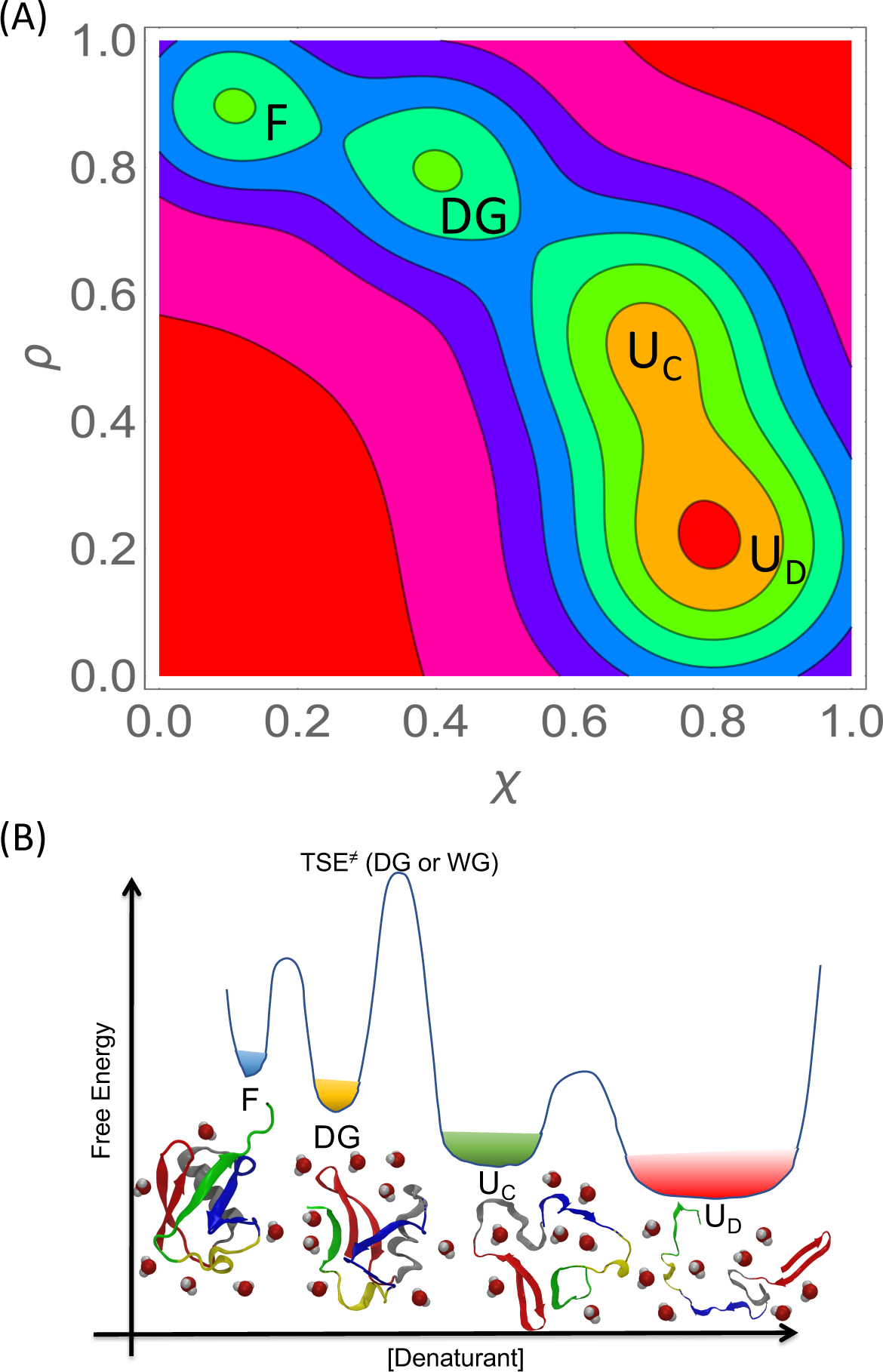
**Box 1 Figure**. (A) Folding landscape of a globular protein projected onto the order parameters, packing density (*ρ*) and the structural overlap function (*ξ* [11]). The folding landscape shows the dry globule state (*DG*) and unfolded collapsed state (*U_C_*) in between the protein folded (*F*) and unfolded states (*U_D_*). As the protein folds from the unfolded state,*ρ* increases and *χ* decreases. The order parameters are described in Box 1. (B) A schematic illustrating the effect of denaturants on the folding landscape. In the folded state, the protein the amino acids are tightly packed and protein hydrophobic core is devoid of water molecules. As the denaturant concentration increases, the folded core loosens forming the molten globule state, which can be wet containing water molecules or dry (*DG* state). As the denaturant concentration increases, the protein unfolds to compact unfolded states (*U_C_*). The barrier between *U_C_* and *U_D_* states is shown only for visual purposes.

The theory intended for describing the coil-globule transition in homopolymers as a function of the solvent quality is altered is strictly valid only when *N* ≫ 1. However, single domain proteins are small (typically contain less than about 200 residues). As a result, the transitions between the states are rounded when studied using *ρ* or *χ*. As a result the values of the order parameters do not change precipitously. In particular, the difference between the radii of gyration between the *U_D_* and *U_C_* states are small [36], requiring accurate measurements over a wide range of [*C*]. The absence of such experiments, until recently, had created a robust and useful controversy. An unambiguous answer to the collapsibility problem, which is important in protein folding and has ramifications for IDPs as well, had therefore remained elusive from an experimental perspective although it has been under theoretical control for twenty five years.

### Simulations and SAXS experiments show that *U_D_* of Ubiquitin undergoes modest compaction

As a case study that illustrates succinctly the many nuances in the collapsibility of the *U_D_* states, we consider the protein Ubiquitin (UB), whose folding has been investigated by changing denaturants [37, 38], mechanical forces [39], and more recently pressure [40]. UB is a 76-residue protein with complicated topology. The crystal structure[41] (PDB ID: 1UBQ) shows that in the folded state it has 5 *β*-strands and 2 *α*-helices (Figure 2A). Experiments [42] and simulations [36, 43] show that the GdmCl midpoint for UB is ∼ 3.8 M at neutral pH (Figure 2B). Simulations [36] in which the effects of GdmCl is modeled using the Molecular Transfer Model [9] show that *R_g_* of the protein increases from ∼ 13 Å to ∼ 25.6 Å as the UB unfolds (Figure 2B). The predicted unfolding transition, as monitored by GdmCl induced swelling, is in excellent agreement with the SAXS experiments [42]. The radius of gyration, 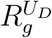, of the *U_D_* state of UB, decreases continuously from ∼ 25.6 Å to ∼ 23 Å as [*GdmCl*] is diluted from 6 to 0.75 M (Figure 2B). The predictions using simulations and the most recent SAXS measurements [44], which show that UB does become compact by 2.2 Å as [*GdmCl*] is diluted from 6 to 0.7 M are in good agreement. Thus, the unfolded states of UB do become compact, albeit only modestly so, as GdmCl concentration is decreased from a high to a low value.

**FIG 2.**
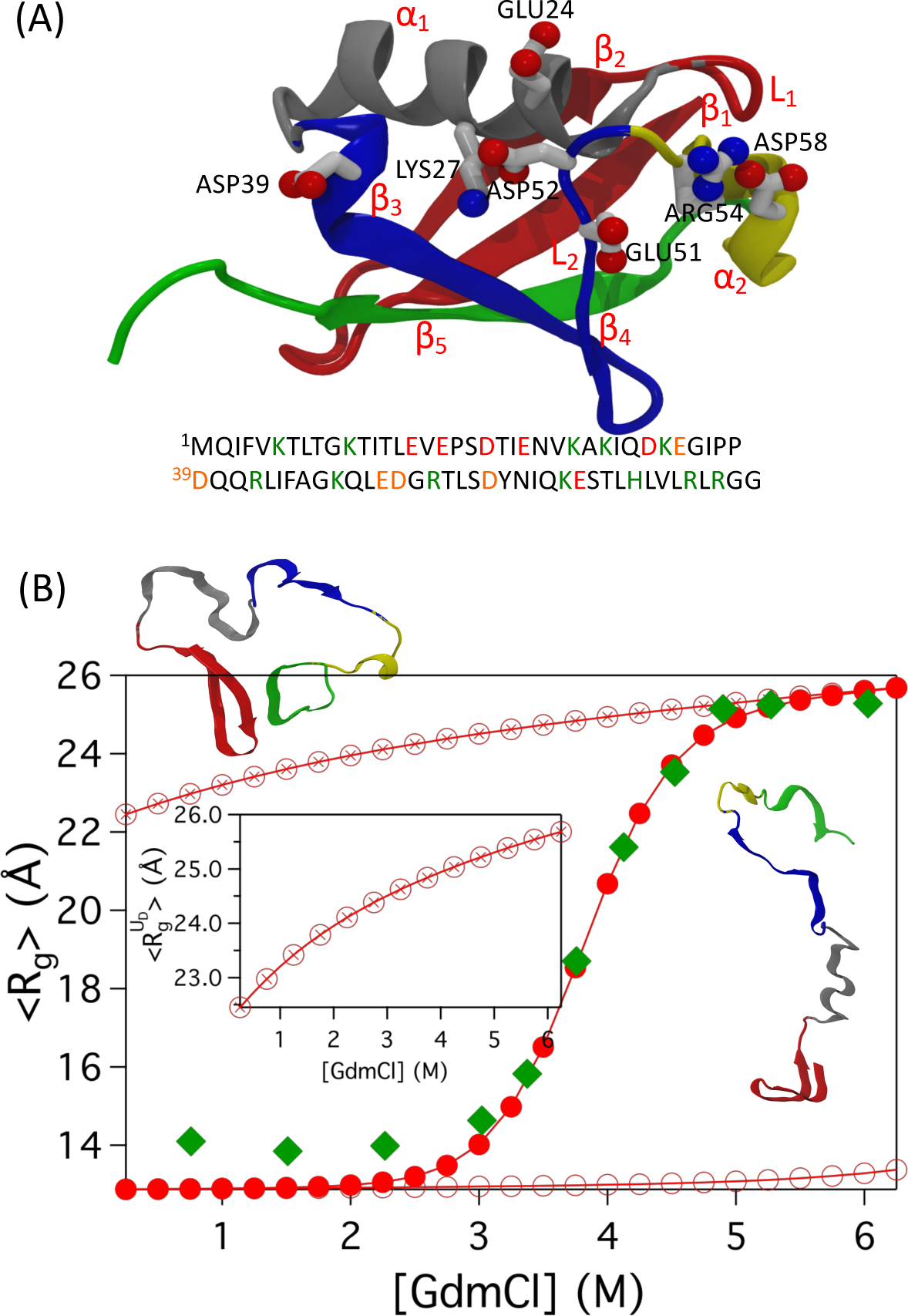
(A) Crystal structure of UB in the folded state (PDB ID: 1UBQ)[41]. UB has 5 *β*-strands (*β*_1_-*β*_5_) shown in red, blue and green, 2 *α*-helices (*α*_1_-*α*_2_) shown in grey and yellow, and it has 2 disordered loops with contacts labeled *L*_1_ and *L*_2_. UB sequence, in a single letter amino acid code, is shown below the structure. Amino acids in green and red letters are positively and negatively charged, respectively. Some of the charged residues are highlighted in the structure. (B) Radius of gyration, *R_g_*, as a function of [*GdmCl*] in neutral pH. Simulation data [36], shown in red solid circles, empty circles and circles with crosses correspond to *R_g_*, 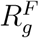 and 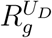, respectively. The inset highlights the continuous decrease in 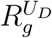 as a function of [*GdmCl*]. Green squares give *R_g_* from SAXS experiments [42]. Simulation snapshots of unfolded UB in the extended and compact form are shown in the figure.

### Is UB in high Urea concentration a random coil?

Simulations [36] and FRET experiments [16, 38] showed that compaction of the *U_D_* state of UB, upon denaturant dilution, is driven by the changes in the solvent quality. To infer the nature of the solvent quality, we calculated the probability distribution of the normalized end-to-end distance (*R_ee_*) of UB, defined as 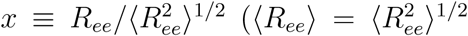 is the mean *R_ee_*). In good solvents, *P*(*x*), should be described by the universal shape corresponding to the self-avoiding walk (SAW) provided *N* ≫ 1. The universal shape of *P*(*x*) for the SAW is given by [45–48]

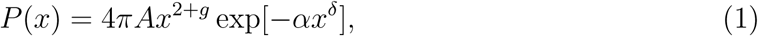

where *v* is the Flory scaling exponent, *g* = (*γ* − 1)/*v*, *δ* = 1/(1 − *v*), and *γ* ≈ 1.1619 for 3 dimensional SAW [49]. The constants, *A* and *α*, are evaluated using the conditions, 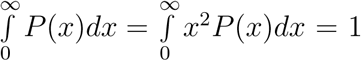.

In acidic pH and high denaturant conditions, [*urea*] = 8 M, *P*(*x*) for UB is well fit using Eq. 1. However, the value of *v* extracted from the fit is 0.75, which is greater than 0.60 expected for long SAWs (Figure 3A). The discrepancy is attributable to the finite size of UB (*N* = 76). In [*urea*] = 2 M, *P*(*x*) shows a bimodal distribution as the C and N termini *β*-strands (*β*_1_ and *β*_5_) (Figure 3A) make contacts with a non-negligible probability leading to a peak in *P*(*x*) at *x* ≈ 0.5 (Figure 3B). This shows that in low denaturant concentrations, the topology of the folded state plays a critical role in determining the extent of compaction of the collapsed states [50].

**FIG 3.**
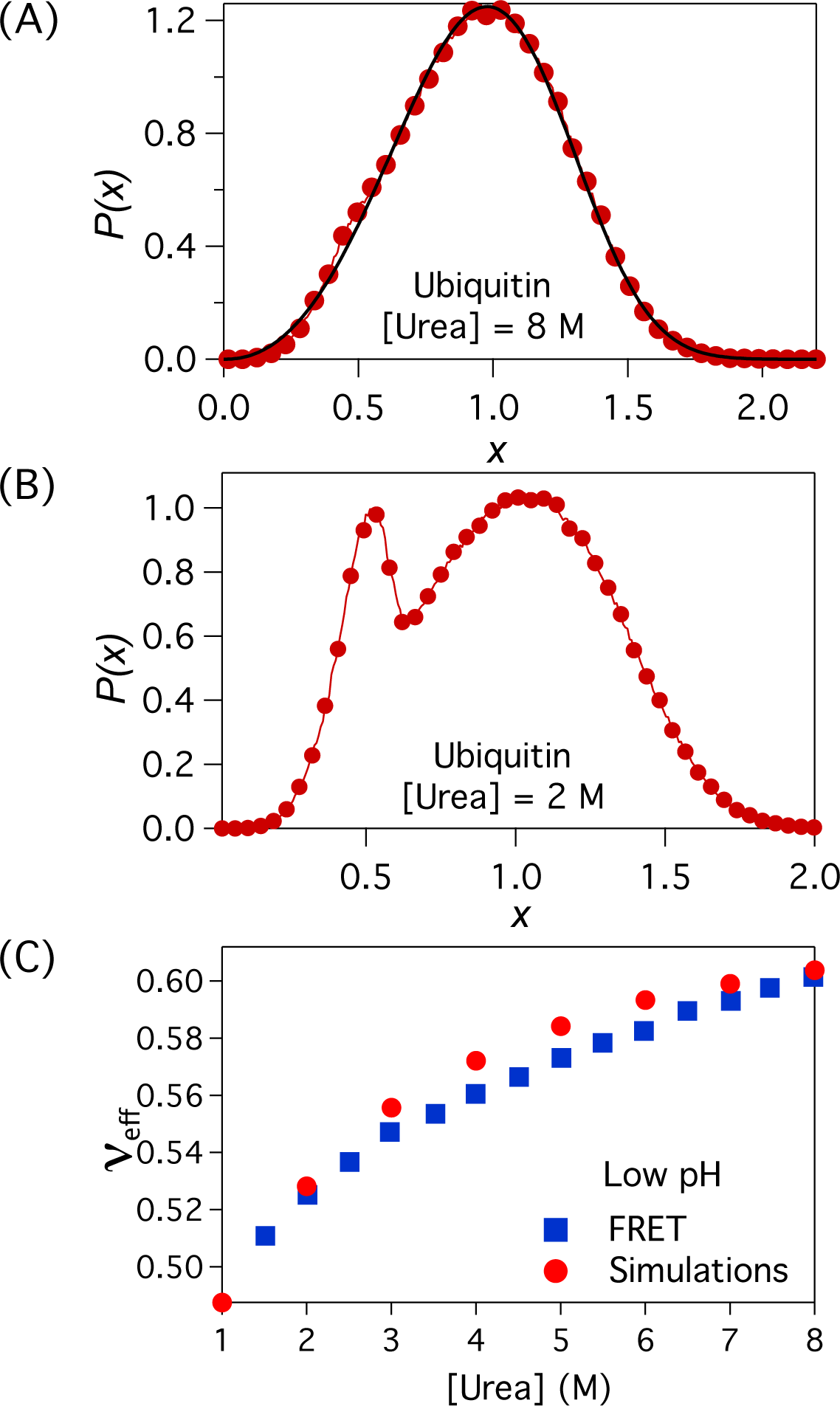
(A) The normalized end-to-end distance, 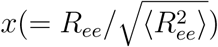, distribution in low pH and [*urea*] = 8.0 M is shown in red circles. The black line is a fit to eq. 1 with *v_eff_* = 0.75. (B) *P*(*x*) in low pH and [*urea*] = 2.0 M. (C) The effective exponent, *v_eff_*, calculated using the relation 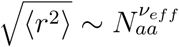, as a function of [*urea*]. Here, 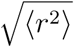 is the intra molecular distance in UB between two residues separated by *N_aa_* residues. The data in blue squares and red circles are from FRET experiments [38] and simulations [36], respectively.

Although the fit of the calculated *P*(*x*) has the form given by Eq. 1, the extracted *v* value from experiments and simulations should be viewed as an effective exponent, *v_eff_*, because there are finite size corrections to the Flory exponent *v* (Box 2). Therefore, based solely on the value of *v_eff_*, we should not conclude that 8 M urea is a good solvent for unfolded UB.

### Solvent quality

The solvent quality for a protein may also be inferred by measuring intramolecular distances between labelled residues, and fitting the results to predictions based on polymer theory. Recently, FRET experiments were performed on UB by positioning the donor and acceptor dyes at different positions to extract the intra chain root mean square distance, 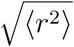 between the dyes[38]. The solvent quality at a particular denaturant concentration is inferred from the effective scaling exponent using the relation, 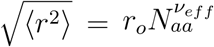, where *N_aa_* is the number of amino acids separating the donor and acceptor dyes (Figure 3C), and *r_o_* is an unknown parameter. The exponent, *v_eff_*, computed using this procedure from both experiments[38] and simulations[36] for ubiquitin shows that *v_eff_* varies continuously from ∼ 0.6 to ∼ 0.5 as [*urea*] is varied from 8 M to 1 M. From this perspective, the solvent quality changes from good solvent like conditions to poor as [*urea*] is diluted (Figure 3C). However, the expectation that for *v* = 0.6, *P*(*x*) must obey the universal shape in eq. 1 is not satisfied because the fit of the simulation data yield *v_eff_* = 0.75. The values of the effective exponent have also been estimated using SAXS data [44] by calculating *R*(*|i − j| ≈ |i − j|^v_off_^*(*R*(*|i − j|* is the mean distance between residues *i* and *j*) from the DSE generated by a new computational method to analyze SAXS data. As outlined in Box 2, this way of estimating *v_off_* is also not without difficulties. Thus, different ways of analyzing the data may not be consistent with each other, casting doubts on the assessment of the solvent quality based on estimates of *v_off_* from FRET or SAXS experiments.

##### Box 2. SAXS and FRET experiments.

The two commonly used methods to measure the radii of gyration (*R_g_s*) of polypeptide chains are SAXS and single molecule FRET experiments. The *R_g_* can be directly calculated using the Guinier approximation to the scattering intensity, *I*(*q*) (Figure 4), which for small *q* is given by 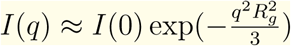 where *I*(0) ∝ to the molecular weight. Thus, the slope of the plot of ln *I*(*q*) versus *q*^2^ yields 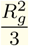. A practical difficulty in determining *R_g_* is that *I*(*q*) has to be measured accurately for values of *qR_g_* ≪ 1. This is particularly important for polypeptide chains for which the changes in *R_g_* are not large. Apparently, the problem is exacerbated at low denaturant concentrations [51], which might contribute to large errors in measuring 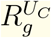.

**FIG 4.**
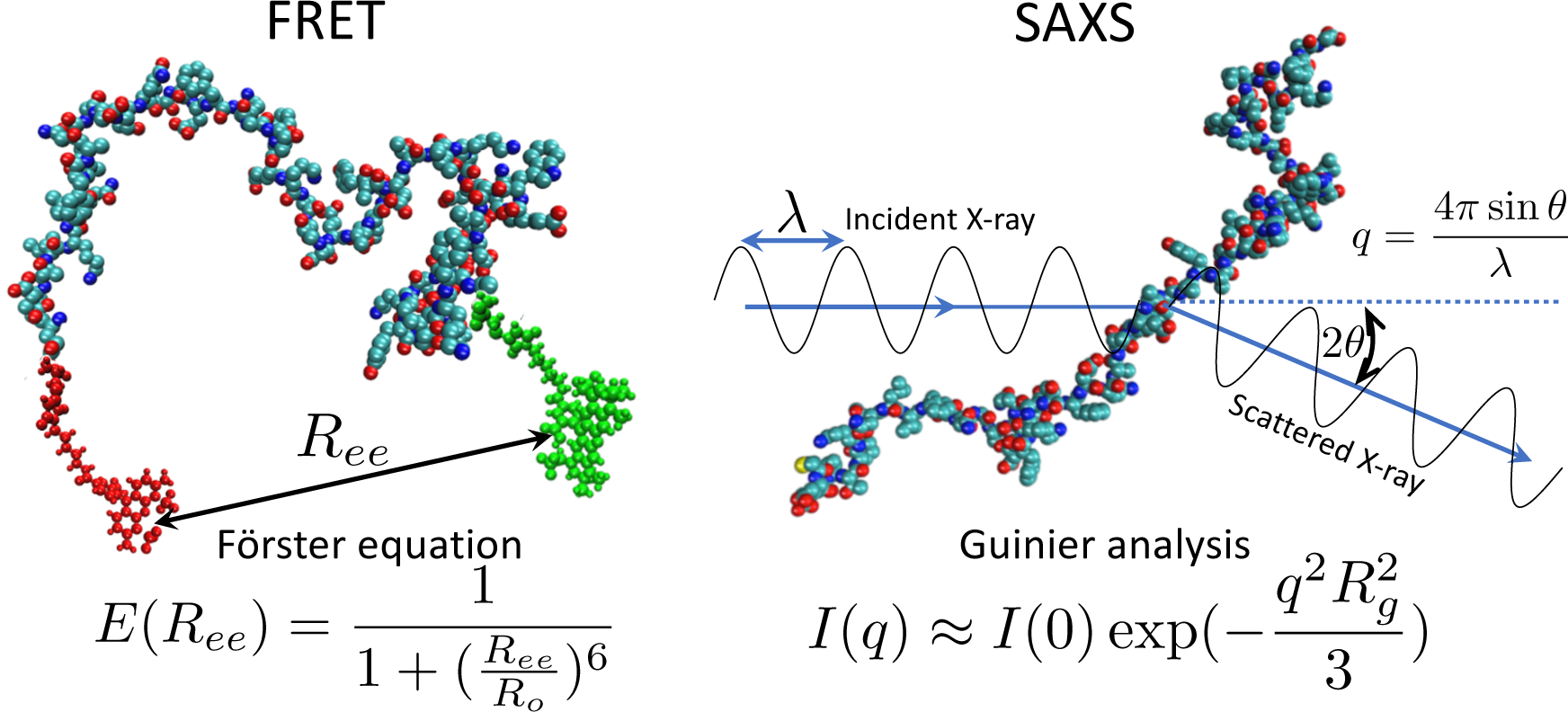
**Box 2 Figure**. Schematic of the FRET and SAXS experiments. In the FRET experiments, the donor and acceptor dyes, shown in red and green, respectively are attached to the protein termini. The efficiency of energy transfer, *E*, between the dyes, which depends on the denaturant concentration is measured. From the measured *E*, the distance between the dyes and the size of the protein is inferred using the Förster relation. In the SAXS experiments, *R_g_* is extracted using the Gunier approximation, which depends on the ratio of the intensities of the incident, *I*(0), and scattered, *I*(*q*), of X-rays at small scattering angles, *θ*.

SAXS also provides information about conformations of the DSE through the distance distribution function, *P*(*r*) given by,

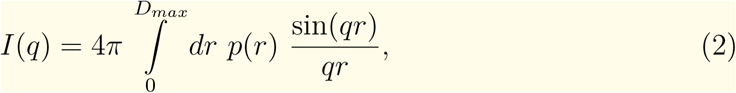

where because of the finite size of the polypeptide chain the upper limit is *D_max_*, which is related to *q_min_*, the smallest wave vector accessible in SAXS experiments. The average value of the square of the radius of gyration is the second moment of *P*(*r*). There are uncertainties in the estimate of *D_max_*, which has to be chosen with care in order to ensure consistency between Guinier approximation and *R_g_* calculated from *P*(*r*). These problems have to be taken into account when SAXS data for the DSE are analyzed.

In smFRET experiments, donor and acceptor dyes are attached to two positions (Figure 4), typically but not always, to the ends of the polypeptide chain. If the dyes are at the ends, then the mean FRET efficiency 〈E〉 is given by,

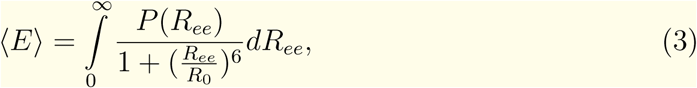

where *P*(*R_ee_*) is the normalized distribution of the end-to-end distance, *R_ee_*, and *R*_0_ is the dye-dependent Forster radius at which *〈E〉* = 0.5. There are limitations in inferring *R_g_* from the measured values of *〈E〉*. (*i*) Calculating *P*(*R_ee_*) using the above equation is a non-trivial inverse problem. A commonly used assumption is that *P*(*R_ee_*) is a Gaussian,

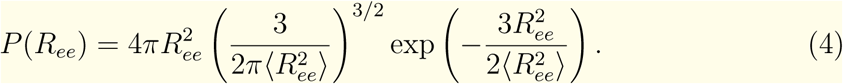

Using eq. 3 and 4, the average radius of gyration of the protein, *〈R_g_〉* is computed using the relation [52], 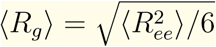, which holds for a Gaussian polymer chain. (ii) The assumption that the unfolded states of polypeptide chains behave as ideal polymers is not accurate, which matters in resolving the apparent SAXS-FRET controversy [53] because in the transition from *U_D_* → *U_C_* the changes in the radius of gyration are small. A consequence is that *〈R_g_〉* of proteins in the denatured ensemble inferred from FRET experiments do not agree with the values obtained from SAXS experiments, and the disagreement is significant for proteins like ubiquitin and protein L [10, 17, 24, 42]. (iii) It is important to point out that for the standard polymer models (Gaussian, Self-Avoiding walks, and Worm-like Chain Model), for which analytic expressions for *P*(*R_ee_*) are available [53], it can be shown that the values of *〈R_ee_〉* exceeds *〈R_g_〉*. This implies that [*δR_g_*]_*FRET*_ > [*δR_g_*]_*SAXS*_ where 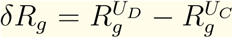. Consequently, FRET experiments exaggerate the extent of compaction of polypeptide chains [9, 36, 43] as the denaturant concentration is decreased. (iv) Finally, the attached dyes could have an effect on the conformations of the polypeptide chains [44], although practitioners of FRET experiments insist that this is not the case [54, 55].

The use of the expression in Eq. 1, with *v* as an adjustable parameter to solve Eq. 3 is also unsatisfactory from a theoretical perspective. The lack of theory, connecting the distance distribution function *P*(*r*) (eq. 2) to *P*(*R_ee_*) for any polymer model other than the Gaussian chain makes it difficult to compare data from the SAXS and FRET techniques in a straightforward manner.

The quality of the solvent (discussed in Box 1) is assessed using *v_eff_* extracted from experimental data. In FRET experiments, *v_eff_* is calculated using [*C*]-dependent *R_g_* = *r_o_N^v_eff_^* where the prefactor *r*_0_ is assumed to be independent of [*C*]. In the most recent SAXS experiments [56] the MFF method is used to generate DSE from which the value of *v_eff_* is extracted using *〈R*(*|i − j|)〉 ≈ |i − j|^v_eff_^* (*〈R(|i − j|)〉* is the mean distance between residues *i* and *j*). The need to know *r_o_* in FRET data analysis is not satisfactory as is the reliance on the accuracy of the DSE conformations generated by the MFF method [44]. If Eq. 1 holds then *〈R(|i − j|)〉 ≈ |i − j|^v_eff_^* calculated for a particular value of *i* and *j* for one protein ought to be identical to the value for another protein as long as *|i − j|* is the same. In addition, *P*(*R(|i − j|*)) should also have the same universal form given by eq. 1 at least when *|i − j|* is large.

In addition, there are finite size corrections [57] to the exponent in the relation, 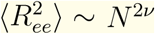, given by,

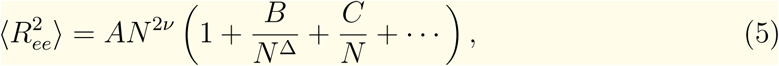

where *v* and the correction to scaling Δ are universal, while *A*, *B* and *C* are system-specific constants. In order to accurately infer the solvent quality using *v*, the corrections to the scaling relation should also be extracted carefully along with *v*, to account for finite *N* [58–60]. Some of these shortcomings together with the broad range in the denaturant concentration range over which the solvent quality changes, render such analyses to be of limited value.

### Extent of compaction is small

The relative compaction in the protein dimensions upon denaturant dilution is small unlike in long synthetic polymers that undergo coil-globule transition, which is akin to a genuine phase transition [61]. The finite-size of proteins is one contributing factor [11, 12]. The relative compaction, Δ, defined as

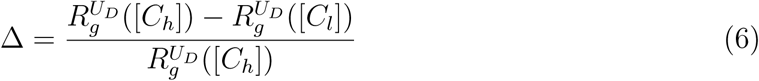

is typically less than 0.2 for a majority of proteins for which reliable simulation and experimental data are available (Table I)[36]. Table I shows that the length variation of the proteins used in experiments to study collapse in the denatured ensemble is small. As a result the change in 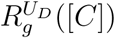 as a function of [*C*] for most of the proteins is not large but is measurable. From Table I it is also clear that there is only a weak correlation between the size of the protein alone and relative compaction. The maximum value of relative compaction is observed for the cold shock protein, which predominantly has a *β*-sheet secondary structure. Thus, besides *N* the topology of the folded protein should also dictate the extent of compaction in the protein unfolded ensemble[50].

**TABLE I:** Relative changes in the radius of gyration of various proteins in the unfolded states between high and low denaturant concentrations

| Protein | $R_g^{UD}([C_h])^c$<br>([GdmCl] = 4-7 M) | $R_g^{UD}([C_l])^d$<br>([GdmCl] = 0-1 M) | $\Delta = \frac{R_g^{UD}([C_h]) - R_g^{UD}([C_l])}{R_g^{UD}([C_h])}$ |
| --- | --- | --- | --- |
| Protein L[73] ($N_{res}=64$) | 26.3 Å ([C] = 7 M) | 22.5 Å ([C] = 0.25 M) | 14.4% |
| Monellin ($N_{res}=96$; Neutral pH)[74] | 27.8 Å ([C] = 7 M) | 25.5 Å ([C] = 0.25 M) | 8.3% |
| PDZ2 Domain[75] ($N_{res}=94$)<sup>a</sup> | 32.2 Å ([C] = 7 M) | 29.8 Å ([C] = 0.25 M) | 7.5% |
| Ubiquitin ($N_{res}=76$; Neutral pH) | 25.9 Å ([C] = 7 M) | 22.5 Å ([C] = 0.25 M) | 13.1% |
| Ubiquitin ($N_{res}=76$; Low pH) | 30.4 Å ([C] = 7 M) | 23.3 Å ([C] = 0.25 M) | 23.4% |
| SH3[76] ($N_{res}=56$) | 23.7 Å ([C] = 7 M) | 20.3 Å ([C] = 0.2 M) | 14.3% |
| Cold Shock[9] ($N_{res}=70$) | 26.4 Å ([C] = 7 M) | 17.8 Å ([C] = 1 M) | 32.6% |
| ACTR[51] ($N_{res}=73$)<sup>b</sup> | 29.7 Å ([C] = 7 M) | 24.7 Å ([C] = 0.3 M) | 16.8% |
| R17d[51] ($N_{res}=116$)<sup>b</sup> | 40.3 Å ([C] = 7 M) | 33.2 Å ([C] = 0.6 M) | 17.6% |
| N49[71] ($N_{res}=38$)<sup>a,b,e</sup> | 16.9 Å ([C] = 6 M) | 15.9 Å ([C] = 0 M) | 5.9% |
| NUS[71] ($N_{res}=82$)<sup>a,b,e</sup> | 31.3 Å ([C] = 6 M) | 24.9 Å ([C] = 0 M) | 20.4% |
| NUL[71] ($N_{res}=114$)<sup>a,b,e</sup> | 35 Å ([C] = 6 M) | 30 Å ([C] = 0 M) | 14.3% |
| PNt[44] ($N_{res}=334$)<sup>b</sup> | 62 Å ([C] = 4 M) | 51.3 Å ([C] = 0.15 M) | 17.3% |
| IBB[71] ($N_{res}=99$)<sup>a,b,e</sup> | 31.2 Å ([C] = 6 M) | 32 Å ([C] = 0 M) | $\approx 0\%$ |
| NLS[71] ($N_{res}=46$)<sup>a,b,e,f</sup> | 23.3 Å ([C] = 6 M) | 24 Å ([C] = 0 M) | $\approx 0\%$ |

<sup>a</sup> Denaturant used is urea.
<sup>b</sup> Data is from experiments.
<sup>c</sup> $R_g^{UD}([C_h])$ is the $R_g$ in the $U_D$ state at high denaturant concentrations.
<sup>d</sup> $R_g^{UD}([C_l])$ is the $R_g$ in the $U_C$ state at low denaturant concentrations.
<sup>e</sup> Sequence details of the IDPs are in the supplementary information of Ref. [71].
<sup>f</sup> The IDP NLS has high mean charge and low mean hydrophobicity. A different buffer, compared to other IDPs, was used in the measurements [71].

### Status of the SAXS-FRET controversy

The details in Box 2 give a glimpse of the difficulties in probing collapsibility of proteins using experiments, which in part explains the SAXS-FRET controversy. Until last year, the persistent claim based on analyses of SAXS data was that the dimensions of the *U_D_* state remains unchanged at all denaturant concentrations including [*C*] = 0 [10, 14]. In an important development, a new analysis method, referred to as Molecular Form Factor (MFF), was used to generate the ensemble of conformations of the unfolded states that are consistent with the measured SAXS profiles. Using MFF and new data for the denatured state ensembles, it was concluded that [44] the radii of gyration of two IDPs decreases between (20-28)% as the polypeptide chains are transferred from 6 M aqueous GdmCl solution to water. For Ubiquitin, 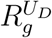 decreases by about 2.2 Å upon a similar change in the GdmCl concentration. In addition, the effective Flory exponent decreases from 0.6 to about 0.5. The analyses of smFRET data for a number of proteins and IDPs consistently indicate that the *U_D_* states become compact. It is gratifying that both camps are in qualitative agreement that the radius of gyration of the *U_C_* states are less than in the *U_D_* states. A perusal of the two commentaries [54, 55] and the response [56] to the article [44] shows that the debate now focusses on what is the quality of aqueous denaturant solution for denatured states of globular proteins and IDPs. We contend based on general theoretical grounds that it is difficult to answer this question using SAXS or FRET. Some of the reasons are outlined in Box 2.

For the subtle issue of collapsibility of the *U_D_* state resolution usually requires theory and accurate simulations that do not require inputs from SAXS or FRET experiments. We believe that the theoretical and simulation results, summarized in Figures 2-5 go a long away in solving the SAXS-FRET controversy. Based on the advances in recent experimental [44, 51] and simulations [36] and theory [50], we summarize the current status of polypeptide chain collapse.

**FIG 5.**
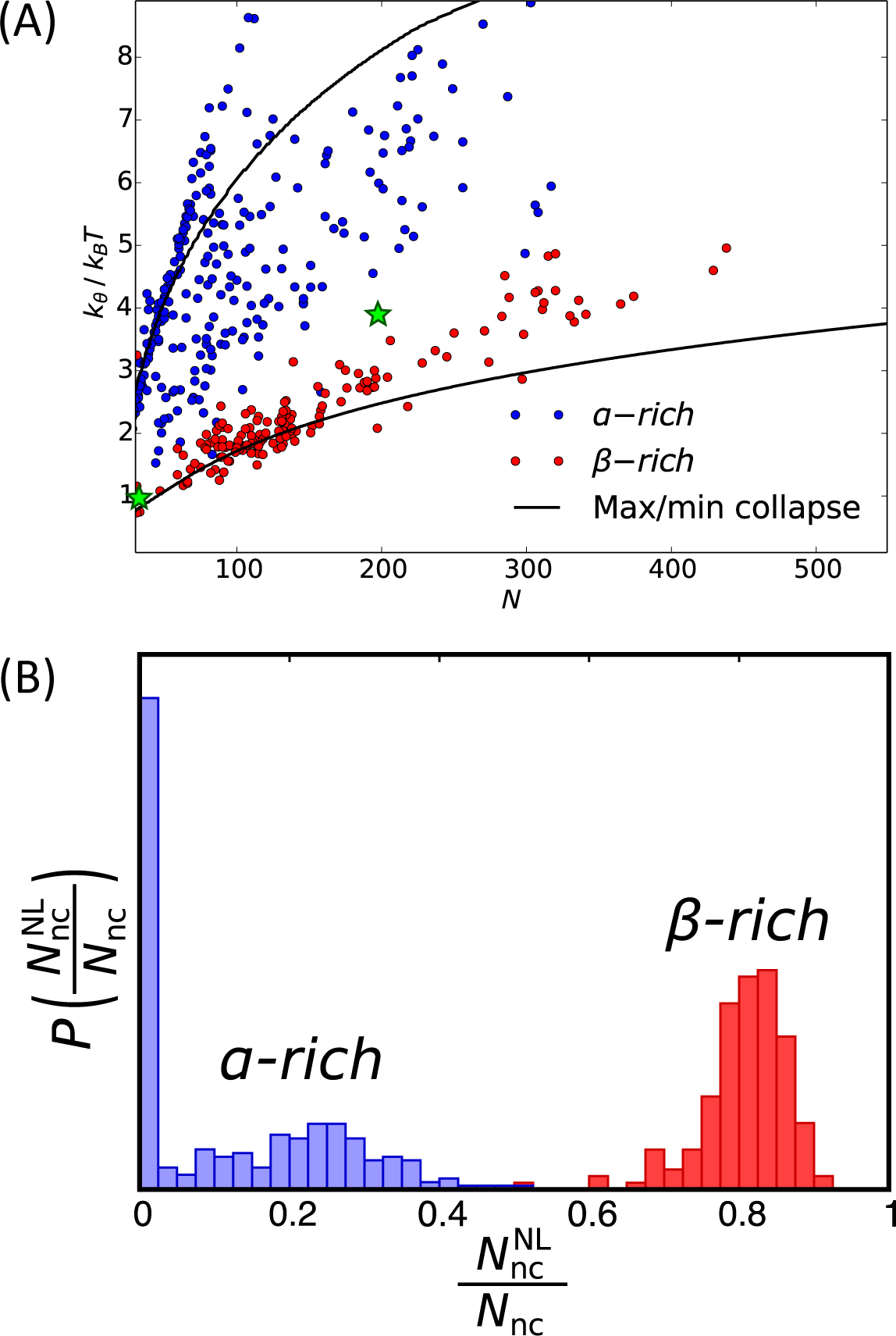
(A) Plot of *κ_θ_*, specifying the average interaction strength between two residues at the *θ*-point in units of *k_B_T*, as a function of the number of residues, *N*. For identical values of *N*, proteins rich in *β*-sheet are more collapsible (have smaller *κ_θ_* values) compared to those rich in *α*-helices. The green star on the left is for a RNA psueudoknot and the other is for the *Azoarcus* ribozyme. (B) 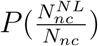 is the distribution of 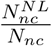, where *N_nc_* is the total number of native contacts in a protein, and 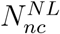 is the number of long range native contacts. Proteins rich in *β*-sheets have more long-range contacts compared to proteins rich in *α*-helices.

- The propensity of unfolded states of globular proteins to collapse is universal [50]. This conclusions is in harmony with analyses of FRET data. Similarly, simulations [36] and SAXS experiments [56] agree regarding the extent of collapse of the *U_D_* state of UB. For example, on decreasing [*GdmCl*] from 4.7 M to 0.9 M, SAXS experiments suggest that 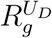 of the *U_D_* state of UB decreases by about 7% whereas our simulations show that, at neutral pH, the decrease is ≈ 13% as [*GdmCl*] decreases from 7 M to 0.25 M (see Table I). The predicted changes are in qualitative agreement with SAXS experiments. We pass the baton to experimentalists to measure changes in 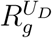 at acidic pH, which we predict is larger (Table I).
- Extraction of the effective Flory exponent from SAXS or FRET data is not straightforward, thus making claims about solvent quality dubious. Strictly speaking, the estimates of the Flory exponent is only meaningful for *N* ≫ 1, which is not satisfied in the studies of single domain globular proteins or even the larger IDPs [56]. Thus, a careful finite size corrections (Eq. 5) must be considered in analyzing data. Nevertheless, theory [12] predicts that for most unfolded states of globular proteins water behaves as a *θ*-solvent, which means that intra protein and water-protein interactions nearly (but not perfectly) cancel with each other. The difficulties in extracting *v_eff_* not withstanding (discussed in Box 2), the cross over in the values of *v_eff_* from good (*v_eff_* ≈ 0.6) to *θ*-solvent (*v_eff_* = 0.5) condition occurs over a broad range of [*C*].
- From Table I, containing the extent of compaction for the denatured states of several proteins, leads to the unexpected conclusion that the length of the protein is not the determining factor in the extent of compaction. It is statistically more closely related to the topology of the folded state, as shown in Figure. 5.

##### Box 3. Theory for Collapsibility

The theoretical basis for assessing collapsibility of a given sequence starts with the Hamiltonian:

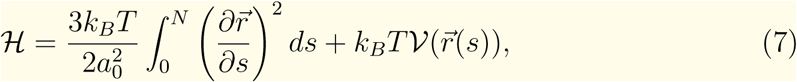

where 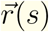 is the position of monomer *s*, and *a*_0_ is the monomer size. The first term accounts for the polymeric nature of polypeptide chains. The interactions between the residues in the above equation are contained in 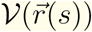 as follows:

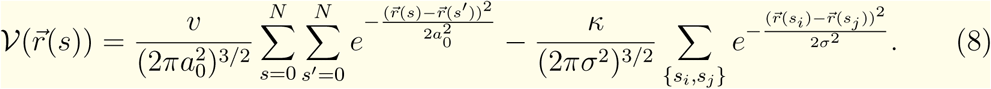

The excluded volume interactions between the residues are represented by the *v* > 0 term. The attraction (term ∝ κ) exists between specific residues, where the sum is over the set of specific interactions between residue pair {*s_i_, s_j_*}. We use the proteins contact maps, computed using the protein data bank (PDB) structures in order to assign the specific interactions [50]. A contact is assigned to any two residues *s_i_* and *s_j_*, if the distance between their *C*_α_ atoms is less than *R_c_* = 8 Å and |*s_i_* − *s_j_*| > 2. For the excluded volume repulsion, the range is the size of the monomer *a*_0_ = 3.8 Å, and for the specific attraction, the range is equal to the average distance between *C*_α_ atoms involved in contact formation, which averaged across a selection of proteins from PBD, is *σ* = 6.3 Å.

Changing the value of *κ*, and hence the strength of attraction, results in the transition between the extended and compact states. Decreasing is analogous to alteration of the concentration of the denaturants ([*C*]). At high [*C*] (*κ* ≈ 0, good solvent) the excluded volume repulsion dominates, while at low denaturant (high *κ*, poor solvent) the attractive interactions are important. The point where attraction balances repulsion is the *θ*-point, and the strength of attraction is *κ_θ_* at the *θ*-point. At the *θ*-point, which is well defined if *N* ≫ 1, the polypeptide chain behaves like an ideal polymer. We define collapsibility using a measure of how easy it is to reach the *θ*-point. In others words, the smaller the value of *κ_θ_* is easier it is for the polypeptide chain to undergo compaction as [*C*] is decreased. Determination of *κ_θ_* is complicated [50] but the final expression may be written in a manner that can be evaluated numerically for any globular protein. The theory makes two major predictions. (1) Polypeptide chain compaction is universal. However, the values of *κ_θ_* depend on the number of residues, *N*, in the polypeptide chains. We showed that [50] *κ_θ_* scales as *N^β^* with *β* > 0. This implies that larger proteins tend to be less collapsible. (2) There is a caveat in the conclusion stated above. The *κ_θ_* values also depend on the folded structures of the globular proteins, with *β*-sheet proteins being more collapsible than *α*-helical proteins.

### *β*-sheet proteins are more collapsible than *α*-helical proteins

The theory outlined in Box 3 allows us to calculate *κ_θ_* for globular proteins. The values of *κ_θ_* (small values imply ease of collapse), plotted in Figure 5 for 2306 proteins, shows that for predominantly *α*-helix proteins (*>* 90%), *κ_θ_* increases rapidly with *N* with most of the proteins lying closer to the minimum collapsibility line. In contrast, the *κ_θ_* values for proteins with high content of *β*-sheet (*>* 70%) are closer to the minimum collapsible line. The values of *κ_θ_* for the two sets are very distinct with minimal overlap. These results show that the extent of collapse of proteins that are mostly *α*-helical is much less than those with predominantly *β*-sheet structures. As the denaturation concentration is lowered below the midpoint, the *U_D_* states reach one of the MECS, which are stabilized predominantly by native-like interactions. Consequently, the extent of compaction at low [*C*] is determined by the topology of the native state, which partly explains the results in Figure 5. Thus, both *N* and even more importantly the topology of the *F* state determine *κ_θ_*.

The reason for ease of collapsibility of proteins that are rich in *β* sheets is that the folded states of these proteins are stabilized by larger number of non-local contacts compared *α*-helical proteins. Indeed, there is a clear separation in the distribution, 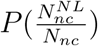 where 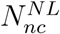 is the number of non-local contacts and *N_nc_* is the total number contacts in a folded protein) between these two classes of proteins. By surveying 2306 proteins, we find that a remarkable separation in 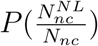 between *α*-rich and *β*-rich proteins, explaining the ease of compaction in the latter compared to the former.

### Compaction as a sequence selection mechanism?

There are two reasons that support the assertion that natural sequences of foldable proteins have evolved to be collapsible. First, precise studies using lattice models show that the constraint of compaction and low energy reduces the number of viable protein structures [3, 7]. Second, the results of our recent theory show that the protein collapse is encoded in the structure [50] to which globular proteins are biased at low denaturant concentrations. Therefore, we conclude that simple biophysical constraint of compaction serves as a plausible selection mechanism in the evolution of foldable sequences. Interestingly, a similar conclusion has been reached for the evolution of viral single stranded RNA molecules [8]. In this context, it has been shown that viral ssRNAs are more compact than random RNA sequences with the same length and similar chemical composition. Thus, based on the principle of parsimony we would suggest that compaction alone might explain evolvability of protein and RNA sequences.

If compaction is a selection mechanism in the evolution of protein sequences then it follows that if the *κ_θ_* values are large, as is the case when *N* increases (*κ_θ_* ≈ *N^β^* with *β* greater than unity), then such proteins might split into multi-domains even at the expense of creating an interface. Each of the individual domains would have *κ_θ_* in a reasonable range, on the order of few *k_B_T* (Fig. 5).

### What about IDPs?

It is estimated that between (30 − 40)% of eukaryotic proteome are either are intrinsically disordered or contain intrinsically disordered regions [62, 63]. Their roles in phase separation resulting in membraneless organelles containing droplets of IDPs [64, 65] has raised the need to understand their statistical properties in isolation. Because they do not form ordered structures they are ideal model systems for obtaining insights into *U_D_* states of globular proteins. Typically, IDPs are low-complexity sequences containing more than usual fraction of charged and polar residues. Statistically they could exhibit properties that are reminiscent of synthetic polyampholytes (PAs) [66] or polyelectrolytes (PEs) [67]. Indeed, FRET experiments [68] on a highly charge IDP have been interpreted using PA theory. Because of the charged nature of certain IDPs salt effects, which are almost always present in the buffer, makes it difficult to analyze SAXS and FRET data. Indeed, the phase diagrams of IDPs in terms of the denaturants and salt concentration are only recently being revealed using theoretical calculations [69, 70]. At a fixed salt concentration, SAXS experiments have shown that the radius of gyration of the 334-residue PNt (an IDP) decreases by about 17% as GdmCl concentration is lowered from about 4M to a low value [44]. Similarly, FRET experiments on both the C and N termini highly charged prothymosin α apparently swell as GdmCl concentration is increased[68]. These findings are in line with the effect of denaturants on the unfolded states of globular proteins. There are exceptions to the general of denaturant-induced expansion of IDPs (see the last two entries in Table I). A complete picture for IDPs will emerge only by simultaneously changing both the denaturant and salt concentrations.

### Concluding Remarks and Remaining problems

For nearly two decades there was no consensus among experimentalists [10, 15], despite early theoretical predictions [11], on whether the sizes of the unfolded states decrease as the concentration of denaturants decreases. Here, we have made a compelling case using the recent advances in theory, simulations, and new SAXS and FRET data that the dimensions of the unfolded states of globular proteins must decrease as the denaturant concentration is lowered, thus putting to bed the long standing SAXS-FRET controversy. The apparent controversy did bring to sharper focus the issue of compaction of the *U_D_* states, which is fundamentally important in understanding the assembly mechanism of proteins. Thanks to advances, just in the last three years [44, 51], we can assert that the compaction of the unfolded states of proteins is universal [50]. However, the changes in the denaturant dependent dimensions in the *U_D_* → *U_C_* is small, thus requiring precise measurements. The debate now seems to have shifted to determining whether water is a good or a *θ*-solvent [54, 56]. For reasons given here, this question can only be answered by measuring the second viral coefficient of the *U_D_* state as function of denaturant concentration.

Despite the overall consensus that collapse is encoded in the evolved protein sequences, there are a few issues that require quantitative answers. Some of these are: (i) There is an urgent need to develop reliable force-fields, which includes the effects of commonly used denaturants, for use in atomically detailed molecular dynamics simulations so that simulations that are independent of experiments can be performed. (ii) In the same vein, a completely model-independent way of analyzing the integral equation connecting FRET efficiency and the distance distribution is needed. (iii) The accuracy of the commonly used Guinier approximation to extract the radius of gyration has been criticized [51, 54] because of substantial errors at low values of the scattering vector (see the figure in Box 2), which is exacerbated at small denaturant concentrations. (iv) The perpetual question of whether dyes attached to polypeptide chains affect their conformations [56] may be partially resolved by doing SAXS and FRET experiments on many other proteins, as has been done recently [71].

There are also exciting questions that deserve scrutiny. First, are aqueous denaturant solutions good solvents for Intrinsically Disordered Proteins? Although there are tentative answers to this question, the analyses methods could be criticized for reasons alluded to in this article. Second, can one quantitatively explain the emergence of multi-domain proteins using collapsibility as an essential physical constraint? We have given a preliminary answer to this question [50] but it remains a speculation. Finally, do these ideas and concepts carry over to ribozymes many of which are also compact? In the RNA field the importance of being compact, driven by cations, has long been accepted especially for ribozymes [72]. However, quantitative description of basic issues such as the dependence of persistence length of RNA and DNA, which is related to flexibility and the propensity of these biological molecules to collapse, is still lacking.

## Acknowledgements

We are grateful to Ben Schuler and Tobin Sosnick for useful discussions and for sharing their data in tabular form. This work was supported by a grant from the National Science Foundation (CHE 16-36424). Additional support from the Collie-Welch Chair through the Welch Foundation (F-0019) is greatly appreciated. GR acknowledges support from the Department of Science and Technology through the grant EMR/2016/001356.

